# Interferon-γ-producing CD4^+^ T cells drive monocyte activation in the bone marrow during experimental *Leishmania donovani* infection

**DOI:** 10.1101/2021.04.26.441484

**Authors:** Audrey Romano, Najmeeyah Brown, Helen Ashwin, Johannes S.P. Doehl, Jonathan Hamp, Mohamed Osman, Nidhi Dey, Gulab Fatima Rani, Tiago Rodrigues Ferreira, Paul M. Kaye

**Affiliations:** York Biomedical Research Institute, Hull York Medical School, University of York, York, U.K; BILHI Genetics, 13016 Marseille, France; Vector Molecular Biology Unit, Laboratory of Malaria and Vector Research, National Institute of Allergy and Infectious Diseases, National Institutes of Health, 12735 Twinbrook Parkway, Rockville, MD 20852, USA; Kennedy Institute of Rheumatology, University of Oxford, Oxford, United Kingdom; Laboratory of Parasitic Diseases, National Institute of Allergy and Infectious Diseases, National Institutes of Health, Bethesda, MD, USA

**Keywords:** visceral leishmaniasis, mouse models, monocytes, CD4^+^ T cells, interferon-γ, IL-10, bone marrow

## Abstract

Ly6C^hi^ inflammatory monocytes develop in the bone marrow and migrate to the site of infection during inflammation. Upon recruitment, Ly6C^hi^ monocytes can differentiate into dendritic cells or macrophages. According to the tissue environment they can also acquire different functions. Several studies have described pre-activation of Ly6C^hi^ monocytes in the bone marrow during parasitic infection, but whether this process occurs during experimental visceral leishmaniasis and, if so, the mechanisms contributing to their activation are yet to be established. In wild type C57BL/6 (B6) mice infected with *Leishmania donovani*, the number of bone marrow Ly6C^hi^ monocytes increased over time. Ly6C^hi^ monocytes displayed a highly activated phenotype from 28 days to 5 months post infection (p.i), with >90% expressing MHCII and >20% expressing iNOS. In comparison, in B6.*Rag2*^-/-^ mice <10% of bone marrow monocytes were MHCII^+^ at day 28 p.i., an activation deficiency that was reversed by adoptive transfer of CD4^+^ T cells. Depletion of CD4^+^ T cells in B6 mice and the use of mixed bone marrow chimeras further indicated that monocyte activation was driven by IFN-γ produced by CD4^+^ T cells. In B6.*Il10*^-/-^ mice, *L. donovani* infection induced a faster but transient activation of bone marrow monocytes, which correlated with the magnitude of CD4^+^ T cell production of IFN-γ and resolution of the infection. Under all of the above conditions, monocyte activation was associated with greater control of parasite load in the bone marrow. Through reinfection studies in B6.*Il10*^-/-^ mice and drug (AmBisome) treatment of B6 mice, we also show the dependence of monocyte activation on parasite load. In summary, these data demonstrate that during *L. donovani* infection, Ly6C^hi^ monocytes are primed in the bone marrow in a process driven by CD4^+^ T cells and whereby IFN-γ promotes and IL-10 limits monocyte activation and that the presence of parasites/parasite antigen plays a crucial role in maintaining bone marrow monocyte activation.

## INTRODUCTION

The bone marrow (BM) is the major site for hematopoiesis in adult mammals, producing all major cell lineages from a pool of committed precursors. Ly6C^hi^ monocytes are derived from a common myeloid progenitor, through intermediates that include monocyte-dendritic cell progenitors and granulocye-monocyte progenitors [1], and with cell fate specified by the transcription factors PU.1, IRF1 and Klf4 as well as by the action of key hematopoietic growth factors such as CSF-1 [2]. Ly6C^hi^ monocytes are released into the bloodstream and migrate to peripheral organs in normal and inflammatory conditions where they contribute to a wide range of physiological and pathological processes including innate and adaptive immune responses, tissue remodeling and tissue repair (reviewed in [3]).

Ly6C^hi^ monocytes are highly plastic and depending on the environment, they can differentiate into a variety of cells including macrophages and dendritic cells (DCs) or maintain a monocyte phenotype [4, 5]. It is unclear, however, whether such terminal differentiation only occurs once in the peripheral tissues or is initiated earlier during BM residency. For example, specialized monocyte progenitors have been demonstrated in the BM and reflect later polarization of monocyte function in the periphery [6]. In addition, Ly6C^hi^ monocytes have the capacity to respond to microbial stimuli during their development in the BM before their egress into the peripheral circulation and ttheir tissue-specific functions are in part pre-programmed [7]. Specifically, infection with *Toxoplasma gondii* led to the secretion of interferon-γ (IFNγ) by NK cells and this cytokine was responsible for monocyte priming and the development of regulatory capacity [7, 8]. These and other studies [1, 6, 9, 10] collectively suggest that Ly6C^hi^ monocytes may become functionally committed prior to their arrival at sites of tissue inflammation or infection.

Although it is known that Ly6C^hi^ monocytes play an important role against various intracellular parasites, the extent to which they contribute to the immune response against different species of *Leishmania* is still unclear, with the data often seemingly contradictory. During *L. major* infection, Ly6C^hi^ monocytes have been demonstrated to have a dual role, on the one hand aggravating during primary infection and on the other hand being protective during secondary infection [11]. This latter function reflects their ability to facilitate rapid recruitment of CD4^+^ T cells at the secondary site of infection and hence to enhance parasite elimination. Others have demonstrated, however, that monocyte-derived DCs are essential for priming protective Th1 response in *L. major* infected mice [12]. The situation is further complicated in models of visceral leishmaniasis, where parasites accumulate predominantly in spleen, liver and BM [13]. Monocytes accumulate in the spleen throughout the course of *L. donovani* infection and play a role in tissue remodeling [14]. Several studies have shown that in absence of Ly6C^hi^ monocytes, B6.*Ccr2*^-/-^ mice fail to generate an effective early Th1 response allowing for a rise in tissue parasite burden load. More recently, a pathogenic role for Ly6C^hi^ monocytes in promoting parasite survival was demonstrated, a finding supported by the observation that emergency hematopoiesis during *L. donovani* infection leads to the differentiation of Ly6C^hi^ monocytes into regulatory monocytes in the BM that contribute to parasite survival [15]. Given that the BM acts as a site of infection, T cells are also recruited in high numbers and we have previously demonstrated that TNF-dependent CD4^+^ IFNγ+ T cells accumulate in significant numbers in the BM resulting in progressive hematopoietic stem cell exhaustion [16] and erosion of the erythropoietic niche [17]. However, the impact of these highly pathogenic CD4^+^ T cells on local monocyte activation has not previously been determined.

In this study, therefore, we used a combination of gene targeted mice, mixed chimeras, antibody deletion and drug treatment to explore the mechanisms underpinning BM Ly6C^hi^ monocyte activation during *L. donovani* infection, and uncover its relationship to BM CD4^+^ T cells, the production of the cytokines IFNγ and IL-10, and parasite load.

## MATERIALS AND METHODS

### Ethic Statement

All animal experiments were carried out in accordance with the Animals and Scientific Procedures Act 1986, under UK Home Office Licence (project licence number PPL 60/4377 approved by the University of York Animal Welfare and Ethics Review Board).

### Animals and infection

B6.CD45.1 (B6.*Ptprc*^a^), B6.CD45.2 (B6.*Ptprc*^b^) and B6.CD45.2.*Rag2*^-/-^ mice used in this study were bred and maintained under specific-pathogen free (SPF) conditions at the Biological Services Facility, University of York. BM cells from mice lacking the *Ifngr1* gene (B6.*Ifngr1*^-/-^) on a B6 background were obtained from the Jackson Laboratory. All mice were between 5–8 weeks of age at the start of experimental work. Mice were infected via the lateral tail vein with 3×10^7^ amastigotes of the Ethiopian strain of *L. donovani* (LV9). To assess the impact of drug-induced parasite clearance, mice were treated once with 10 mg/kg Amphotericin B (AmBisome®) at day 28 post-infection and killed 72 hours later. Animals were killed by CO2 asphyxia and cervical dislocation at the time points specified. Spleen and liver parasite burden was expressed as Leishman-Donovan units (LDU), where LDU was equal to the number of parasites/1000 host nuclei multiplied by the organ weight in milligrams. BM parasite burden was determined by limiting dilution assay (LDA). Briefly, two-fold serial dilutions in 96-well flat bottom microtiter plates were performed in OMEM medium supplemented with 20% FCS. The plates were scored microscopically for growth and the number of parasites in each tissue was determined from the highest dilution at which parasites could be grown out after 7–14 days incubation at 26°C.

### Cell extraction

The spleen was mechanically disrupted and incubated in RPMI with 0.25 mg/ml Collagenase IV and 0.1 mg/ml DNase 1 for 30min at RT. Spleen tissue was then forced through a 70µm strainer to obtain a single cell suspension. Cell pellets were resuspended in ACK Lysing buffer (5min at RT) then washed with RPMI. BM cells from tibias were flushed with a 26g needle and cold RPMI. To obtain a single cell suspension, cells were run through a 70µm cell strainer and spun down. Cell pellets were resuspended in ACK lysing buffer 5min at RT and washed. Cells were resuspended in RPMI 1640 medium (Life Technologies) supplemented with 10% heat-inactivated FCS, 4 mM L-glutamine, 10 mM HEPES, 100 U/ml penicillin and 100 μg/ml streptomycin and kept on ice. Cells were counted with trypan blue under a light microscope.

### Flow Cytometry and cell sorting

For immunolabelling, cells were washed and labelled with LIVE/DEAD Fixable Dead Cell Stains (Thermo Fisher Scientific) to exclude dead cells. Cells were incubated with anti-Fc III/II (CD16/32) receptor Ab (2.4G2), followed by surface staining with various combination of the following antibodies for 30min at 4°C in the dark: CD11b (M1/70), Ly6G (1A8); Sca1 (D7), CD45 (30-F11), CD45.1 (A20), CD45.2 (104), CD3e (145-2C11 and UCHT1), CD4 (RM4-5 and 4SM95), CD8β (H35-17.2), TCR-β (145-2C11), CD11c (N418), Ly6C (HK1.4), MHC-II (M5/114.15.2), MerTK (2B10C42), CD40 (XX), CD80 (XX), CD86 (GL-1), CD64(X54-5/7.1). Staining for CCR2 employed AlexaFluor 700 anti-CCR2 (475301) and was done prior to surface staining at 37°C for 20-30min.

In some experiments, cells were stimulated with Brefeldin A at 10µg/ml alone or in combination with Phorbol-12-myristate-13-acetate (PMA) (Sigma-Aldrich) and ionomycin (Sigma-Aldrich) for 4h at 37°C, then fixed and permeabilized with the eBioscience™ Intracellular Fixation & Permeabilization Buffer Set according to the manufacturer instructions and stained with IL-10 (JES5-16E3), NOS2 (CXNFT), TNF (MP6-XT22), IFNγ (XMG1.2).

All Abs were from eBiosciences, BD Biosciences, Biolegends or R&D systems. Data were collected using FACSDiva software on BD LSR Fortessa X-20 (BD Biosciences), and analyzed using FlowJo software (TreeStar). Forward-scatter and side-scatter width was employed to exclude cell doublets from analysis. Cell sorting to > 95% purity was performed on a MoFlo Astrios (Beckman Coulter).

### Mixed bone marrow chimeras and adoptive T cell transfer

BM cells were purified as described previously. B6.CD45.1 recipient mice were irradiated with 850 rad from an X-ray source using a two split does regimen and reconstituted with 10^6^ cells from B6.CD45.1 and B6.*Ifngr1*^-/-^.CD45.2 mice admixed in a 50:50 ratio. Mice were maintained on oral antibiotics for the first 6 weeks post reconstitution. Mice were infected at 7 weeks of chimerism and killed 28 days after the infection. For T cell adoptive transfer, B6.*Rag2*^-/-^ CD45.2 mice infected for 21 days received 10^6^ CD4^+^ or CD8^+^ T cells isolated from the BM of 28 day-infected B6.CD45.1 mice. Mice were killed 7 days post-transfer for analysis.

### Treatment with anti-IL10R, anti-CD4 monoclonal antibody

CD4^+^ T cells were depleted by administering 400 μg of InViVoMAb anti-mouse CD4 (clone GK1.5) intra-peritoneally, twice weekly beginning on day 14 post-infection and for 2 weeks. Control mice were injecting with the same amount and at the same frequency with InVivoMAb rat IgG2b isotype control (anti-keyhole limpet hemocyanin; LTF-2). IL-10 signaling was blocked by injecting 250μg of InVivoMAb anti-mouse IL-10R (clone 1B1.3A; CD210) intraperitoneally every 3 days for 14 days starting on day1 post-infection. InViVoMAb rat IgG1 Isotype control (anti-trinitophenol; TNP6A) was administered to control mice. All antibodies and isotype controls were purchased from BioXcell (Lebanon, USA).

### Statistical analysis

Numbers of animals used in each experimental group is detailed in the figure legends. Statistical analysis was performed using students’ t test with p<0.05 considered significant. All analyses were conducted using GraphPad InStat (version 6) software. Number of repeats are indicated in corresponding figure legends.

## RESULTS

### Inflammatory monocytes are primed in the BM of B6 but not B6.Rag2^-/-^ mice during L. donovani infection

The activation of monocytes in the BM during infection has yet to be explored in the context of *L. donovani* infection. In contrast to infection of immunocompetent B6 mice, infection in B6.*Rag2*^-/-^ mice was characterised by delayed hepato-splenomegaly and unchecked parasite growth in spleen, liver and BM over a 5 month period (**Fig 1A-C**). Inflammatory monocytes (iMo), defined as CD45^+^CD11b^+^Ly6G^-^Ly6C^hi^CCR2^+^ cells (**Fig 1D**), decreased in number early after infection, as noted by others [15], but then transiently increased in number in the BM of B6 mice being ∼2-fold higher in absolute number at 28 day p.i. In contrast in B6.*Rag2*^-/-^ mice, no early decline in absolute monocyte numbers was noted early after infection, but a similar transient ∼2 fold increase was observed at d28 (**Fig 1E**). To determine whether these iMo were found in a primed state, we analysed expression of MHCII and iNOS ([7]) over the course of infection. MHCII expression was low in both strains of mice for the first 14 days p.i. and increased from d28 p.i., with marked differences between B6 and B6.*Rag2*^-/-^ mice (**Fig 1F and G**) In B6 mice, MHCII was abundant on almost all BM monocytes from d28 p.i. onwards whereas in B6.*Rag2*^*-/-*^ mice, only 20% of monocytes expressed MHCII at later time points. This lack of activation in immunodeficient mice was even more evident when measuring intracellular iNOS (**Fig 1H**). These data suggest that iMo are primed in the BM during *L. donovani* infection and that adaptive immunity is required to drive this response. Furthermore, as infection is controlled in B6 mice, the extent of BM monocyte activation as measured by iNOS generation wanes, despite maintenance of high MHCII expression.

**Figure 1:**
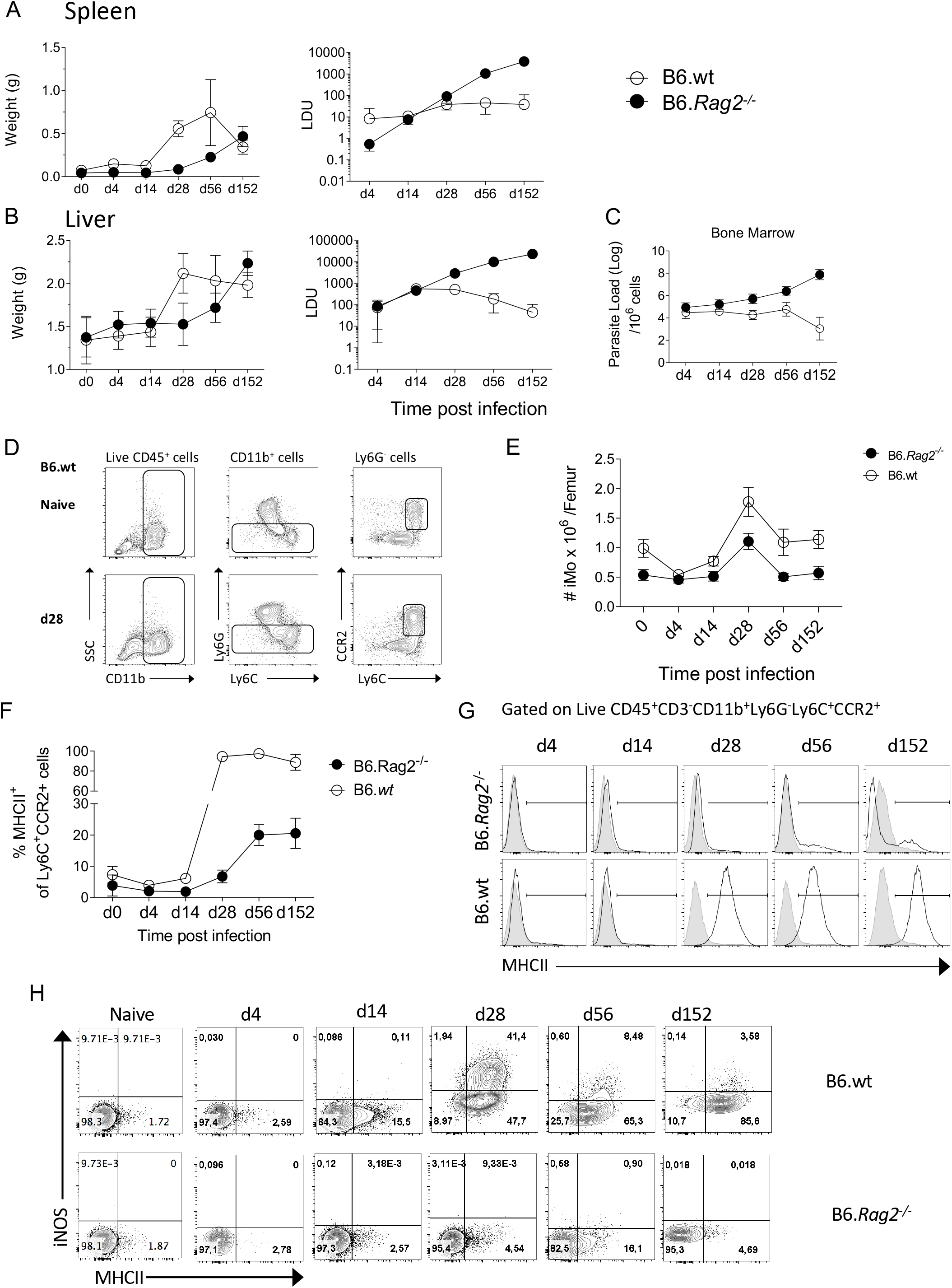
Monocytes are activated during *L. donovani* infection in B6 but not in B6.*Rag2*^*-/-*^ mice. **A and B**. Weight (left panel) and Leishman Donovan Unit (right panel) in the spleen (A) and liver (B) of B6 mice and B6.*Rag2*^*-/-*^ at different time point during the course of the infection. **C**. Parasite load per one million cells in the bone marrow measured by limiting dilution assay. **D**. Gating strategy to identify Ly6C monocytes in BM. **E**. Absolute number of iMo cells per femur. **F and G** MHCII expression on BM Ly6C^+^CCR2^+^ cells shown as percentage (F) and as flow histogram plots (G) in B6 and B6.*Rag2*^*-/-*^ mice during the course of the infection. H) iNOS and MHCII expression on Ly6C^+^CCR2^+^ monocytes. Data are derived from analysis of 8 to 10 individual mice (4 to 5 / experiment) of each strain at each time point pooled from two independent experiments and are shown as mean ± SD. Data in G and H are representative plots.

### CD4^+^ T cells are the main producers of IFNγ and drive BM monocyte activation in an IFNγ-dependent manner

IFNγ has been described as a key cytokine controlling monocyte activation. To identify probable sources of IFN*γ* in the BM of *L. donovani*-infected mice, we stimulated BM leucocytes at different times p.i. with PMA and Ionomycin. Approximately 60-70% of CD4^+^ T cells had the capacity to produce IFNγ at d28 and these cells persisted to 8wks p.i ([16] and **Fig 2A**). In contrast, ∼ 30% of CD8^+^T cells were capable of producing IFNγ at d28 p.i. and this frequency declined thereafter. No IL-10 production was observed by either CD4^+^ or CD8^+^ BM T cells under these stimulation conditions (**Fig 2A**). CD4^+^ T cells represented ∼75% of all IFNγ^+^ cells in the BM of B6 mice at d28 (**Fig 2B**). Of note, the expansion of IFNγ-producing CD4^+^ T cells preceded that of MHCII expression on iMo (**Fig 2C**), suggesting a causal relationship. To address this directly, we used an adoptive transfer model in B6.*Rag2*^-/-^ recipients (**Fig 2D**). Adoptively-transferred CD4^+^ isolated from d28-infected B6 mice retained their capacity to produce IFN*γ* in B6.*Rag2*^-/-^ hosts (**Fig 2E**) and induced MHCII expression on recipient BM iMo (**Fig 2F and G**). In contrast, CD8^+^ T cells produced less IFN*γ* after adoptive transfer than in infected B6 donor mice and poorly induced monocyte activation (**Fig 2E-G**). Importantly, transfer of CD4^+^ T cells into a naïve B6.*Rag2*^-/-^ mice did not lead to BM iMo activation, ruling out an effect due to homeostatic T cell expansion (**Fig S1**) and demonstrating a requirement for cognate antigen recognition in vivo. The origin of CD4^+^ T cells (naïve versus d28 infected B6 mice) did not change the outcome of the experiments (**Fig S1)**.

**Figure 2:**
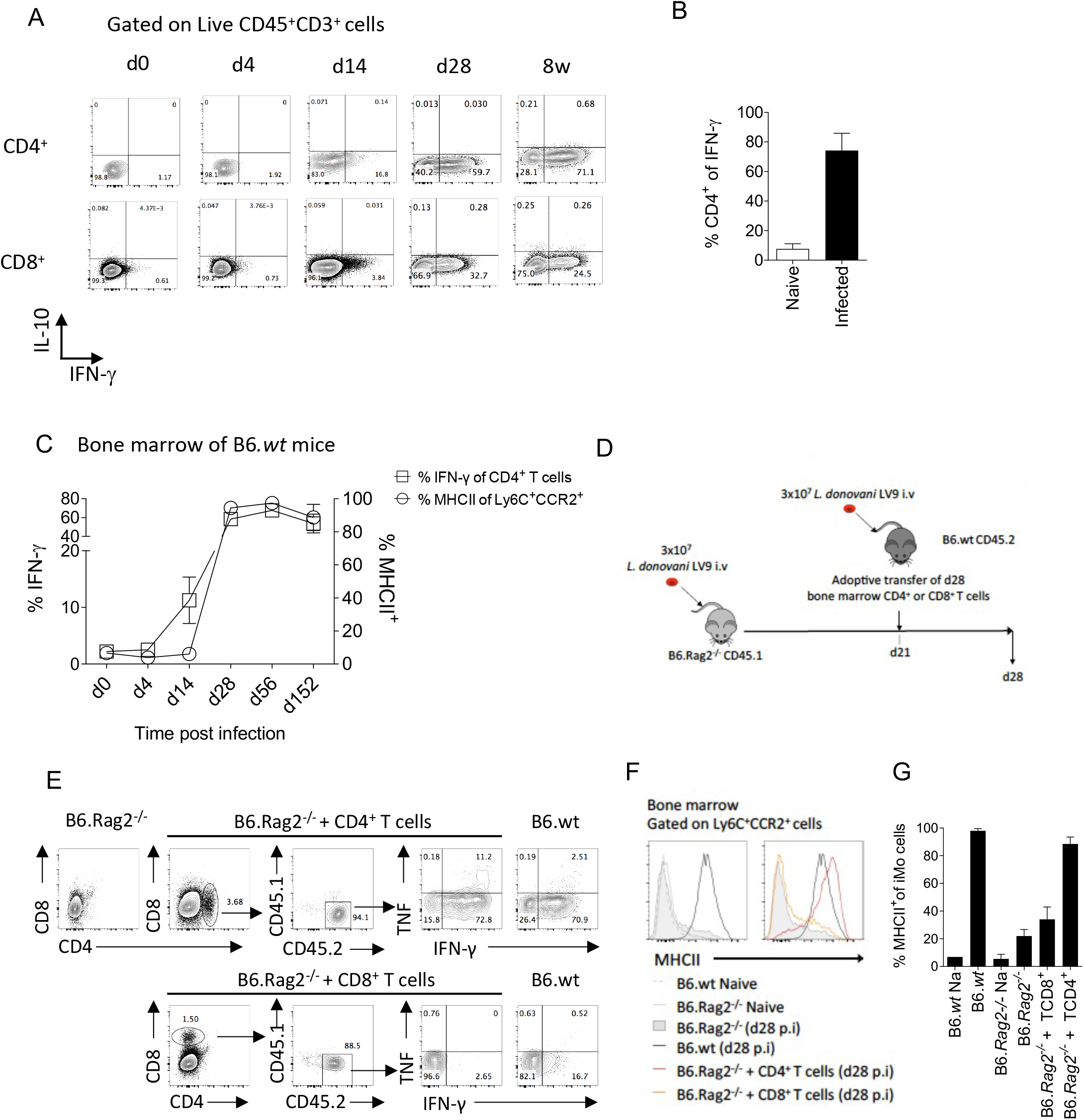
CD4^+^ T cells are the main producers of IFNγ in the bone marrow of B6 mice and contribute to local monocyte activation. **A** Dot plot indicating representative expression of IL-10 and IFNγ by CD3^+^CD4^+^ T cells at different time points p.i. Cells were stimulated with PMA/Ionomycin. **B** Percentage of CD4^+^ T cells expressing IFNγ. **C** IFNγ expression by BM CD4^+^ T cells and MHCII expression on Ly6C^+^CCR2^+^ monocytes. **D**. Protocol for adoptive transfer of T cells. **E**. Cytokine production by adoptively transferred BM CD4^+^ or CD8^+^ T cells 1 week post transfer into recipient *L. donovani*-infected B6.*Rag2*^*-/-*^ mice. Cells were stimulated with PMA/Ionomycin. **F and G** MHCII expression on Ly6C^+^CCR2^+^ monocytes in the BM of the mice one week post adoptive T cell transfer, shown as histogram plots (**F**) and percentage (**G**). Data are derived from analysis of 3 to 4 individual mice of each strain per group and are shown as mean ± SD. Data are pooled from 2 independent experiment. Data in **E** and **F** are representative plots.

To independently confirm the role of CD4^+^ T cells in iMo activation, we used the alternate approach of depleting CD4^+^ T cells from immunocompetent mice. We depleted CD4^+^ cells in infected B6 mice from day 14 p.i to d28 p.i. with anti-CD4 depleting antibody, the time frame over which iMO MHCII expression increases (**Fig 2C**). After CD4^+^ T cell depletion, the frequency of total BM cells producing IFNγ in the BM was reduced by approximately 70% (0.8% vs 2.8% in anti-CD4 treated vs. control mice; **Fig 3A**). Although depletion was incomplete, we observed a reduction in MHCII expression in iMo reflected in both the percentage of MHCII^+^ cells (**Fig 3B and C**) and in median fluorescent intensity of expression (**Fig 3D**). Of note, parasite load in the BM of the mice treated with anti-CD4 mAb was increased compared to control mice (**Fig 3E**), indicating that the reduction in CD4^+^ T cells and presumably IFNγ production had a significant impact on host resistance to *L. donovani*. This is supported by the observation of a significant decrease of parasite number in the BM of adoptively transferred B6.*Rag2*^*-/-*^ mice as compared to their control (**Fig S2**).

**Figure 3:**
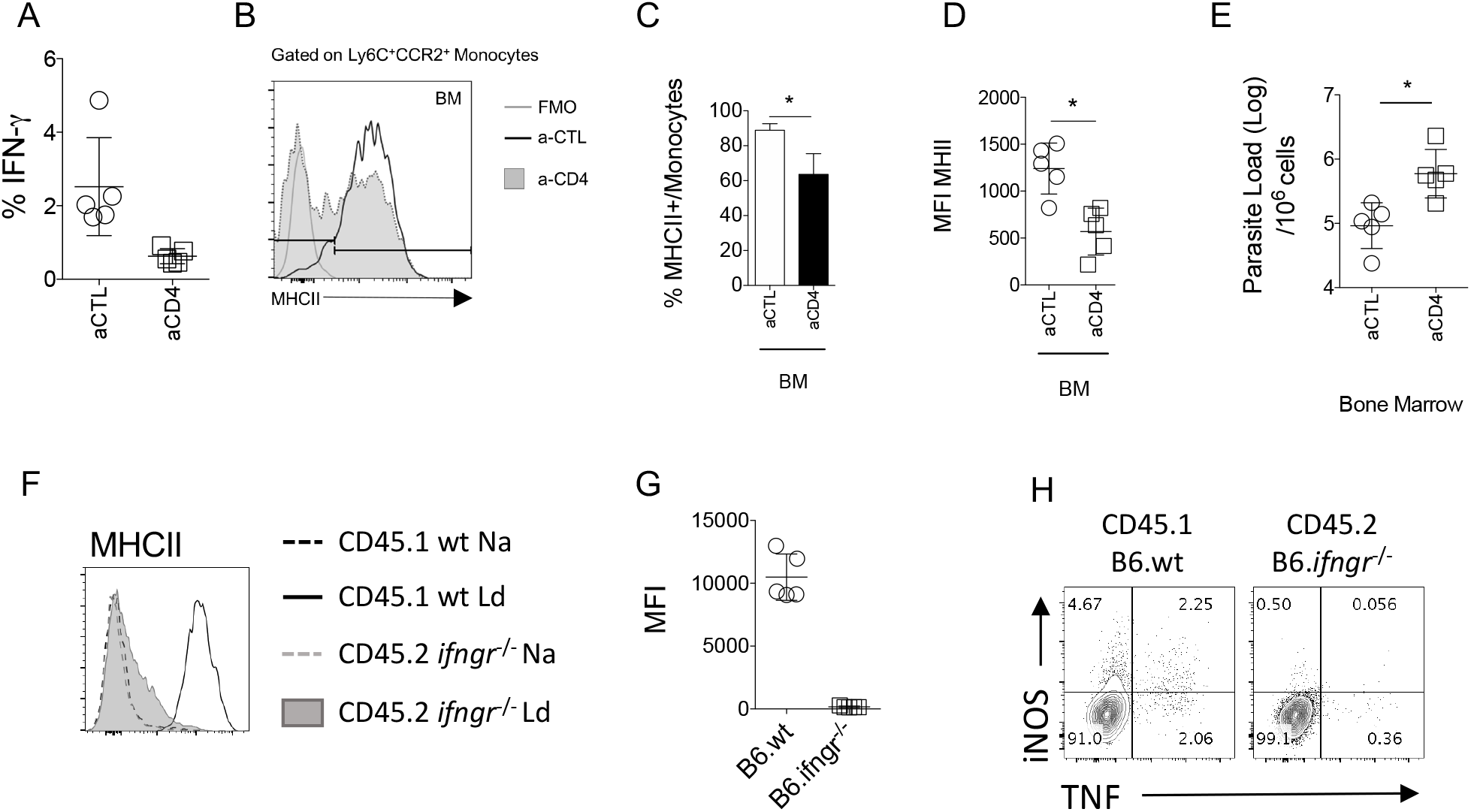
Monocytes respond to IFN-γ produced by CD4^+^ T. **A-E**) At day 14 p.i, B6.wt mice were injected with an anti-CD4 depleting antibody or a control antibody for 14 days. All the data shown correspond to BM results. **A**) Total percentage of IFNγ produced in the BM. Histogram and bar graph representing the expression (**B**) and percentage (**C**) respectively, of MHCII on Ly6C^+^CCR2^+^ monocytes. **D**) MFI of MHCII and CD80 on the Ly6C^+^CCR2^+^ monocytes. **E**) Parasite burden measured in infected B6 at d28 by limiting dilution assay. **F and G**) Naïve lethally irradiated B6.CD45.1 recipient mice received a 50:50 of BM cells from B6.CD45.1 and B6.*Ifngr1*^*-/-*^ CD45.2 mice. Subsequently infected with 3×10^7^ *L. donovani* amastigotes for 28 days. **F**) At day 28, the level of activation of Ly6C^+^CCR2^+^ monocytes cells was determined based on the level of expression of MHCII. **G**) Dot plot representative of the production of NOS2 and TNF by CD45.1 or CD45.2 Ly6C^+^CCR2^+^ monocytes in the BM. Data are the pool of 5 mice/group/experiment. Mean +/-SD is shown. Experiment was performed once.

Finally, in order to verify whether BM iMo respond directly to IFNγ, we used a mixed chimera model. Irradiated B6.CD45.1 mice were reconstituted with B6.CD45.1 and B6.*Ifngr1*^-/-^.CD45.2 BM cells in a 1:1 ratio (**Fig S3A)**. At day 28 p.i, CD45.1^+^ (wild type) but not CD45.2^+^ (*Ifngr1*^-/-^) iMo were primed in the BM, as determined by expression of MHCII (**Fig 3F and G**). *Ifngr1*^-/-^ iMo also produced minimal levels of NOS2 and TNF compared to wild type iMo in these IFNγ-sufficient mixed chimeras (**Fig 3H)**. As expected, *Ifngr1*^-/-^ and wild type CD4^+^ T cells produced equivalent amount of IFNγ in the BM (**Fig S3B**). Collectively, these data indicate that BM CD4^+^ T cells produce IFNγ and that IFNγ signalling on BM iMo is necessary to induce their activation in *L. donovani* infected mice.

### IL-10 inhibits iMo activation and restrains parasite elimination

Having demonstrated the main pathway related to iMO activation in the BM of *L. donovani*-infected mice, we next wished to determine whether this was constrained in any way by regulatory cytokine production. Although IL-10 has been long implicated as a negative regulator of macrophage activation in multiple other settings [18], its role in iMO activation is not known. As BM CD4^+^ T cells did not produce demonstrable IL-10 after in vitro restimulation (**Fig 2A**), we used B6.*Il10*^-/-^ mice to evaluate the possible role of IL-10 in regulating IFNγ-mediated iMO activation. B6.*Il10*^-/-^ mice have previously been shown to be resistant to *L. donovani* infection in terms of liver and spleen parasite loads [19]. In BM, B6.*Il10*^-/-^ mice had reduced parasite load at all time points measured (**Fig 4A**) and at d56 parasites were undetectable in the BM of B6.*Il10*^-/-^ mice when measured by LDA. We then measured iMO activation by MHCII and iNOS expression. At d14 p.i., 18.9% ± 11.2 of iMo in were NOS2^+^ compared to 0.85% ± 1.54 in B6 mice (p=0.005). Similarly, 98.5% ± 6.0 of iMo in B6.*Il10*^-/-^ mice expressed MHCII compared to 42.85% ± 12.73 in B6 mice (p<0.0001). Together, these data indicate a more rapid kinetic for iMo activation in the absence of IL-10 (**Fig 4B-F**).

**Figure 4:**
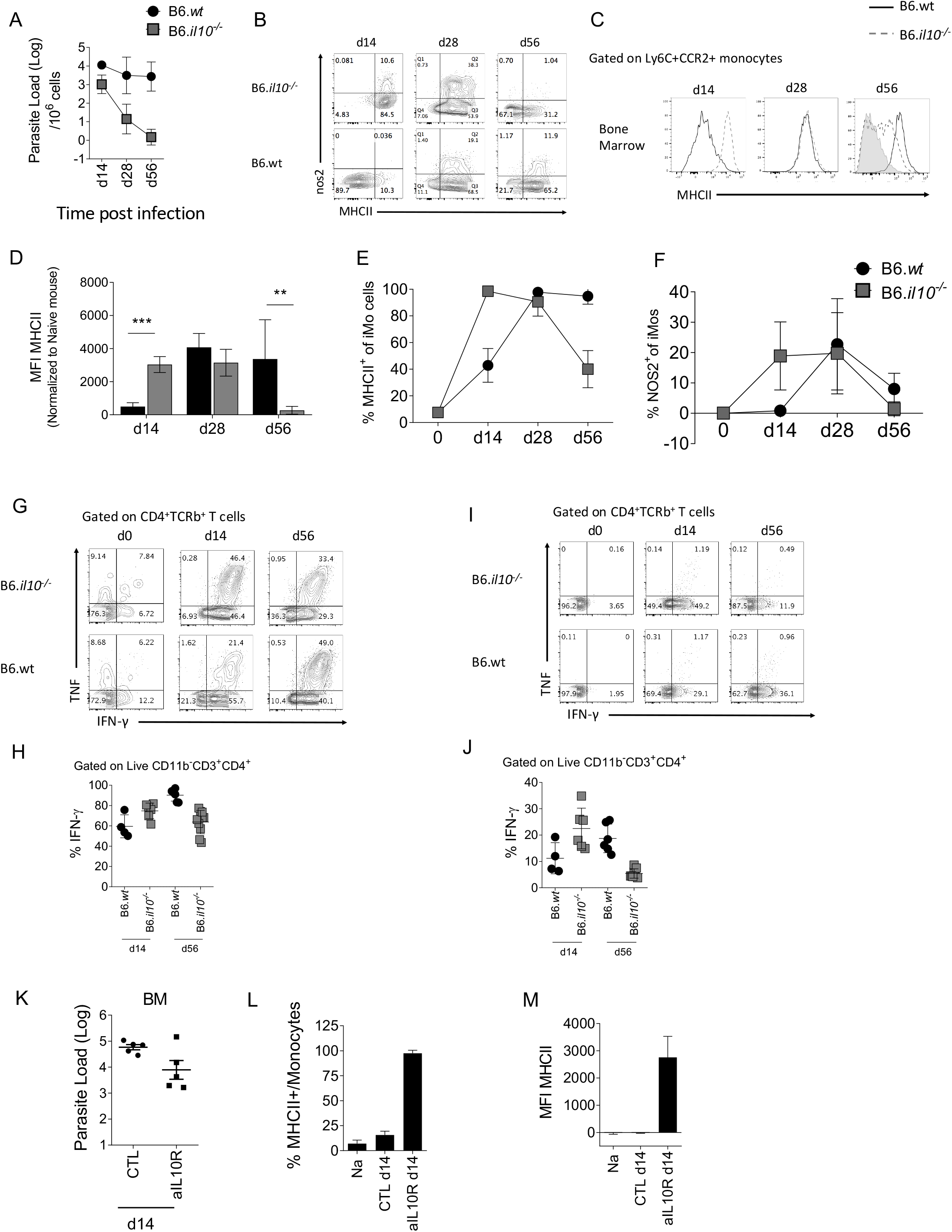
Absence of IL-10 changes the kinetics of monocyte activation in the BM. B6 and B6.*Il10*^*-/-*^ mice were infected with *L*.*donovani* amastioges in the tail vein for 14, 28 and 56 days. Data from the BM are shown **A-G. A**) Parasites burden in the bone marrow determined by limiting dilution assay at the indicated time point. **B**) Dot plot representative of nos2 and MHCII expression by Ly6C^+^CCR2^+^ monocytes. **C-F**) Histogram (**C**), bar graph showing MFI (**D**), percentage of MHCII (**E**) and percentage of nos2 **(F**) on BM Ly6C^+^CCR2^+^ monocytes. **G and I**) Production of IFN-γ and TNF by CD4^+^ T cells in the BM at the different time point post-infection and (**H and J**) percentage of IFN-γ produced by CD4^+^ T cells after PMA-Ionomycine-BFA stimulation for 4h (**G and H**) and *ex vivo* (BFA stimulation only, (**I and J**). **K-M**) B6 mice infected with *L. donovani* were injected with an anti-IL-10 depleting antibody or the isotype control for 14 days starting 1 day prior to the infection **K**) Parasite load determined by limiting dilution assay in the BM at day 14 p.i. **L**) Percentage and **M**) MFI of MHCII expressed on Ly6C^+^CCR2^+^ monocytes. For A-J, data are derived from analysis of 6 to 7 individual mice (2 to 4 / experiment) of each strain at each time point pooled from two independent experiments and are shown as mean ± SD. Data in B, C, G and I are representative plots. For K-M, data are the pool of 5 mice/group/time point. Mean ± SD is shown. Experiment was performed once.

Nevertheless, at d28 p.i., iMO activation for iNOS production was similar in the presence or absence of IL-10 (22.7 ± 15.0 vs. 19.8 ± 13.4, respectively; p=0.7) as was MHCII expression (97.8 ± 2.7 vs. 90.5 ± 10.6; p=0.13). Surprisingly, whereas iMo activation was sustained in wild type mice at d56 p.i., it was reduced at this later time point in IL-10-deficient mice, as determined by frequency of MHCII^+^ cells (94.7 ± 6.0 vs. 40.0 ± 13.9; p<0.0001; **Fig 4E**). In contrast, iNOS production declined somewhat in B6 mice but to a greater extent in IL-10-deficient mice (8.0 ± 5.1 vs. 1.5 ± 5.1% respectively; p=0.02; **Fig 4F**).

As parasites were no longer detectable at d56 p.i in B6.*Il10*^-/-^ mice (**Fig 4A**), we hypothesized that the presence of parasites may be necessary to maintain iMo activation status, providing an antigen depot to allow continued stimulation of IFN-producing CD4^+^ T cells. To test this hypothesis, we measured the level of IFNγ production by CD4^+^ T cells at d14 and d56 p.i in the BM of B6 and B6.*Il10*^-/-^ mice (**Fig 4G - J**). At d14 p.i., CD4^+^ T cells from B6.*Il10*^-/-^ mice produced more IFNγ than those in B6.*Il10*^-/-^ mice after both polyclonal stimulation (**Fig 4G and H**) and more strikingly directly ex vivo (**Fig 4I and J**). In contrast, at d56 p.i., this situation was reversed, again more strikingly with direct ex vivo measurement of IFNγ (**Fig 4I and J**). These data suggest that IL-10 initially restrains iMo activation, but that at later time points, IL-10 may indirectly contribute to sustaining the level of iMo activation via maintenance of parasite load and ongoing effector CD4^+^ T cell IFNγ production.

To confirm whether early IL-10 restrains iMo activation, we used the alternate approach of blocking IL-10 signalling in B6 mice by administration of anti-IL-10 receptor antibody for 14 days, starting at day 1 p.i. (**Fig 4K**). Blockade of IL-10 signalling led to a decrease in parasite load in the BM, as expected from the results observed in B6.*Il10*^-/-^ mice (**Fig 4A**), as well as an increase in iMo activation, as measured by MHCII expression (**Fig 4L and M**).

### The presence of parasites in the BM contributes to iMo activation

To determine whether parasite load / and or antigen availability plays a role in determining the level of iMo activation, we re-infected B6.*Il10*^-/-^ mice at d56 p.i., a time at which they had cleared their primary infection (**Fig 5A**). One day post-re-infection, *Leishmania* parasites were detected in the BM of re-infected mice (**Fig 5B**). Strikingly, BM iMo from these re-infected mice were highly activated, returning to levels of MHCII expression seen at d14 p.i. (**Fig 5C-E**).

**Figure 5:**
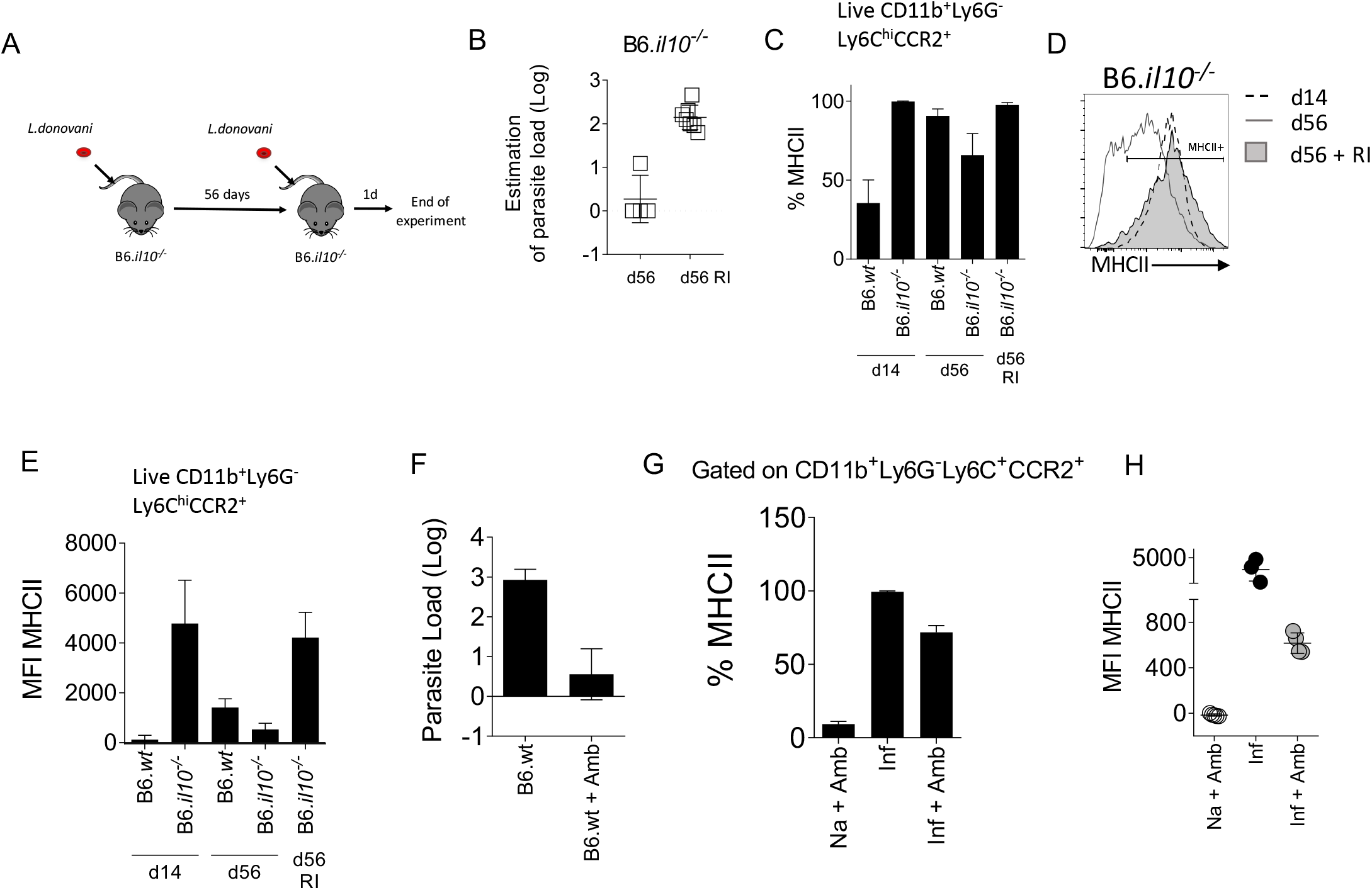
The presence of *L. donovani* parasites in the BM is necessary to maintain monocyte activation. **A-E**) B6.*Il10*^*-/-*^ mice were re-infected (RI) at d56 with 3×10^7^ *L. donovani* amastigotes and killed 1 day post re-infection. **A**) Experimental strategy. **B**) Parasite load per one million cells determined by limiting dilution assay in the BM at day 56 or 1 day post RI. **C**) Percentage of MHCII on Ly6C^+^CCR2^+^ monocytes in the BM. **D**) Level of MHCII expressed by the BM Ly6C^+^CCR2^+^ monocytes represented on histogram. **E**) MFI of MHCII on Ly6C^+^CCR2^+^ monocytes in the BM of the indicated mice. **F-H**) B6 mice infected for 28 days were treated with 10mg/kg of Ambisome®. Data were collected 3 days post-treatment. **F**) Parasite load measured by limiting dilution assay in the BM. Percentage (**F**) and MFI (**H**) of MHCII expressed on Ly6C+CCR2+ monocytes. In A-E, data are from 3 to 5 mice per group / experiment. Experiments were performed twice and pooled data are shown. In F-H, data were derived from 3 to 4 mice / group.

We then conducted the reciprocal experiment, using the anti-leishmanial drug AmBisome® to clear parasite load prematurely in day 28-infected B6 mice. Treatment with Ambisome® for 3 days reduced BM parasite load to less than 10 parasites/10^6^ cells (**Fig 5F**) and the level of activated iMo was reduced significantly, as determined by a reduction in the frequency of MHCII^+^ cells (99.37% ± 0.61 reduced to 71.73% ± 4.63 in control vs drug-treated mice; **Fig 5G**) and a reduction in MFI for MHCII (3619 ± 1360 to 616.3 ± 90.2 in control vs drug-treated mice; **Fig 5H**). Thus, forced early reduction in parasite load is accompanied by reduced iMo activation in IL-10-sufficient B6 mice.

Collectively, these data suggest a model whereby parasite load in the BM is an essential driver of CD4^+^IFNγ^+^ T cell-dependent iMO activation, and that IL-10 serves to regulate this pathway via its capacity to regulate macrophage leishmanicidal activity.

## DISCUSSION

Our data extend a previous study of BM monocytes that also demonstrated an elevation of MHCII expression on Ly6Chi iMo in the BM early during *L. donovani* infection [15] by extending analysis into the chronic period of infection and though analysis of the mechanisms responsible for this finding. Notably, we demonstrate that the changes seen in B6 mice in terms of MHCII expression is a direct consequence of IFNγ-signalling and in turn that this IFNγ is produced predominately by CD4^+^ T cells and opposed by IL-10.

Through the independent experimental approaches of gene targeted mice and mAb blockade, we have provided evidence that IL-10 impacts on the activation status of BM monocytes during *L. donovani* infection. Although IL-10 production by splenic CD25^-^ Foxp3^-^ CD4^+^ T cells has been reported to correlate with disease progression in experimental VL [20, 21], in the BM we found no evidence that CD4^+^ (or CD8^+^) T cells produced appreciably levels of IL-10 over 8 weeks course of infection (Figure 2A). This finding is in accord with the notion that the CD4^+^ T cell population recruited to the BM during infection may be selected for IFNγ production [16]. In the absence of IL-10 production by T cells, other cellular sources are therefore implicated. In spleen and liver, these have been shown to include NK cells [22] and CD11c^hi^ conventional dendritic cells [21] in addition to macrophages and monocytes themselves [23-26]. Furthermore, innate activation of B cells to produce Il-10 has been demonstrated in the context of *L. donovani* infection [27] and plasma cells producing IL-10 have been implicated in directing myeloid cells differentiation under homeostatic conditions [28]. Further investigation with for example IL-10 transcriptional reporter mice will be required to confirm the identity of the IL-10-producing cells in the BM of *L. donovani*-infected mice and their relative roles in regulating monocyte activation.

In this study, we did not aim to directly assess the role of iMo in disease outcome, a subject that has been studied by others. Satoskar and colleagues have previously reported that iMo were associated with enhanced parasite loads in the spleen of B6 mice and that these iMo showed an activation profile biased at the transcriptional level towards arginase production. In contrast, iMo in the liver of these *L. donovani*-infected mice where more biased towards iNOS production [29]. Paradoxically, blockade of iMo ingress to the spleen and liver using a CCR2 antagonist led to improved host resistance in both tissues, suggesting that phenotypic characteristics measured in vitro may not always reflect in vivo function. In contrast, another study using CCR2-deficient mice failed to demonstrate a role for iMo in host resistance and hepatic granuloma formation [30]. We have not directly measured arginase in BM iMO in this study, but iNOS production was clearly elevated throughout the course of infection through a IFNγ-dependent pathway, most likely involving STAT1 signalling [29]. Whilst this study also showed that iMO in the peritoneal cavity could efficiently phagocytose *L. donovani* amatigotes, they did not show that this directly occurred at the major sites of infection. Abidin et al [15] demonstrated that alterations in myelopoiesis led to the generation of regulatory iMO that suppress protective immunity and these monocytes were identified to be parasitised in the BM. Phenotypic changes in MHCII and Ly6C expression suggested exposure to IFNγ, in keeping with our direct demonstration of a role for IFNγR signalling, but also up-regulation of galectin-3, associated with alternate activation [31]. Further studies will be necessary to identify if those changes in monocytes activation affects anti-leishmanial immunity in other sites of infection such as the spleen and the liver.

Our in vivo data also provide an in vivo example of the intimate relationship between immune activation and parasite load. Although IL-10 deficiency leads to a rapid acceleration and augmentation of monocyte activation, this is transient and by d56 post infection, monocyte activation in B6.*Il10*^-/-^ mice has returned to a homeostatic baseline. One explanation for this apparently paradoxical observation is that maintenance of monocyte activation and local IFNγ production is dependent upon the presence of parasites in the BM environment, a suggestion borne out by experimentation. Thus, reduction of parasite load by drug treatment rapidly reduced expression of MHCII on BM monocytes whereas reinfection of previously cured B6.*Il10*^-/-^ mice with *L. donovani* led to a rapid increase in BM monocyte MHCII expression. Similarly, adoptive transfer of CD4^+^ T cells from infected mice into naïve recipient hosts does not lead to monocyte activation, indicating a need for parasites to trigger this response in the BM monocytes. The lack of monocyte activation in the BM of infected B6.*Rag2*^*-/-*^ mice suggests that the parasites alone are not sufficient for such activation. Rather, IFNγ-producing CD4^+^ T cells and *Leishmania* are both required to induce this response. Currently, we cannot distinguish between a model in which parasite antigens contribute to CD4^+^ T cell activation through cognate pathways of antigen presentation or one in which parasites induce bystander CD4^+^ T cell activation through innate (e.g. TLR) signalling pathways.

Alternatively, parasites and or their products may directly stimulate monocytes through similar engagement of pattern recognition receptors, though this itself is insufficient to drive MHCII and iNOS expression in the absence of T cell-derived IFNγ. Nevertheless, our data clearly demonstrate the importance of parasite load as a parameter influencing local immunoregulatory pathways.

Whilst the kinetics of induction of MHCII expression following T cell activation was expected based on the know role of IFNγ in the transcriptional regulation of MHCII expression on myeloid cells [32], the rapid change seen after pathogen clearance with AmBisome® in terms of both the frequency of MHCII^+^ BM monocytes and the abundance of MHCII was surprising. One explanation for a reduced frequency of MHCII^+^ monocytes is that MHCII^hi^ iMo are released from the BM as a consequence of a change in the balance of retention / egress signals [33] and that newly produced monocytes fail to become activated. CXCL12, the ligand for the main retention receptor CXCR4, is produced locally by BM and splenic stromal cells but expression is reduced at the mRNA and protein level in these tissues as well as in the BM following *L. donovani* infection [17, 34-36]. In contrast, the monocyte attracting chemokines CCL2, CCL7 and CCL8 remain highly expressed in spleen [36] and splenic CCL2 mRNA accumulation remains elevated even after AmBisome® treatment [37]. Alternatively, the more marked reduction in MFI for MHCII on BM monocytes is suggestive of cell intrinsic changes in protein expression. E3 ubiquitin ligases of the MARCH family play a role in regulating cell surface expression of a variety of membrane proteins [38]. Of note, MARCH1 plays a central role in IL-10-mediated post translational regulation of surface MHCII expression in human monocytes and other myeloid cells [39, 40], in addition to regulating monocyte differentiation [41]. Further studies would be needed to further dissect this novel aspect of monocyte MHCII expression following parasite clearance.

In summary, we have demonstrated that BM monocyte activation is a prominent feature of *L. donovani* infection in mice and reflects the outcome of an intricate balance between IFNγ, IL-10 and parasite load. Little is currently known about the role of BM monocytes in the progression of human VL and during the response of patients to therapy. The studies outlined here, together with the relative ease and safety with which bone marrow aspirates can be obtained from VL patients, provide a foundation to explore in more detail this aspect of myeloid cell function during human VL.

## CONFLICT OF INTEREST

The authors declare that the research was conducted in the absence of any commercial or financial relationships that could be construed as a potential conflict of interest.

## AUTHOR CONTRIBUTIONS

Conceptualisation: AR, PMK; Data curation: AR, PMK; Formal analysis: AR; Funding acquisition: PMK; Investigation: GA, NB, HA, JSPD, JH, MO, ND, GFR, TRF; Methodology: AR, PMK; Project administration: PMK; Resources: PMK; Supervision: PMK; Visualisation: AR; Writing, original draft: AR, PMK; Writing, review and editing: AR, PMK.

## FUNDING

This work was funded by a Wellcome Trust Senior Investigator Award to PMK (WT104726).

## ACKNOWLEDGMENTS

The authors thank the staff of the Biological Services Facility (BSF) and Biosciences Technology Facility for their support in animal husbandry and flow cytometry, respectively.

## DATA AVAILABILITY STATEMENT

The raw datasets for Figures 1-5 of this study will be posted to OSF on publication.

## Notes

### Competing Interest Statement

The authors have declared no competing interest.

## REFERENCES

1. Yanez A, Coetzee SG, Olsson A, Muench DE, Berman BP, Hazelett DJ, et al. Granulocyte-Monocyte Progenitors and Monocyte-Dendritic Cell Progenitors Independently Produce Functionally Distinct Monocytes. Immunity. 2017;47(5):890–902 e4. Epub 2017/11/23. doi: 10.1016/j.immuni.2017.10.021. PubMed PMID: 29166589; PubMed Central PMCID: PMCPMC5726802.

2. Terry RL, Miller SD. Molecular control of monocyte development. Cell Immunol. 2014;291(1-2):16–21. Epub 2014/04/09. doi: 10.1016/j.cellimm.2014.02.008. PubMed PMID: 24709055; PubMed Central PMCID: PMCPMC4162862.

3. Jakubzick CV, Randolph GJ, Henson PM. Monocyte differentiation and antigen-presenting functions. Nat Rev Immunol. 2017;17(6):349–62. Epub 2017/04/25. doi: 10.1038/nri.2017.28. PubMed PMID: 28436425.

4. Zigmond E, Varol C. Two Roads Diverge in the Sick Liver, Monocytes Travel Both. Immunity. 2020;53(3):479–81. Epub 2020/09/17. doi: 10.1016/j.immuni.2020.08.006. PubMed PMID: 32937148.

5. Zigmond E, Varol C, Farache J, Elmaliah E, Satpathy AT, Friedlander G, et al. Ly6C hi monocytes in the inflamed colon give rise to proinflammatory effector cells and migratory antigen-presenting cells. Immunity. 2012;37(6):1076–90. Epub 2012/12/12. doi: 10.1016/j.immuni.2012.08.026. PubMed PMID: 23219392.

6. Menezes S, Melandri D, Anselmi G, Perchet T, Loschko J, Dubrot J, et al. The Heterogeneity of Ly6C(hi) Monocytes Controls Their Differentiation into iNOS(+) Macrophages or Monocyte-Derived Dendritic Cells. Immunity. 2016;45(6):1205–18. Epub 2016/12/22. doi: 10.1016/j.immuni.2016.12.001. PubMed PMID: 28002729; PubMed Central PMCID: PMCPMC5196026.

7. Askenase MH, Han SJ, Byrd AL, Morais da Fonseca D, Bouladoux N, Wilhelm C, et al. Bone-Marrow-Resident NK Cells Prime Monocytes for Regulatory Function during Infection. Immunity. 2015;42(6):1130–42. Epub 2015/06/14. doi: 10.1016/j.immuni.2015.05.011. PubMed PMID: 26070484; PubMed Central PMCID: PMCPMC4472558.

8. Park J, Hunter CA. The role of macrophages in protective and pathological responses to Toxoplasma gondii. Parasite Immunol. 2020;42(7):e12712. Epub 2020/03/19. doi: 10.1111/pim.12712. PubMed PMID: 32187690.

9. Serbina NV, Pamer EG. Monocyte emigration from bone marrow during bacterial infection requires signals mediated by chemokine receptor CCR2. Nat Immunol. 2006;7(3):311–7. Epub 2006/02/08. doi: 10.1038/ni1309. PubMed PMID: 16462739.

10. 2 Zhu J, Chen H, Huang X, Jiang S, Yang Y. Ly6C(hi) monocytes regulate T cell responses in viral hepatitis. JCI Insight. 2016;1(17):e89880. Epub 2016/10/26. doi: 10.1172/jci.insight.89880. PubMed PMID: 27777980; PubMed Central PMCID: PMCPMC5070953.

11. Romano A, Carneiro MBH, Doria NA, Roma EH, Ribeiro-Gomes FL, Inbar E, et al. Divergent roles for Ly6C+CCR2+CX3CR1+ inflammatory monocytes during primary or secondary infection of the skin with the intra-phagosomal pathogen Leishmania major. PLoS Pathog. 2017;13(6):e1006479. Epub 2017/07/01. doi: 10.1371/journal.ppat.1006479. PubMed PMID: 28666021; PubMed Central PMCID: PMCPMC5509374.

12. Leon B, Lopez-Bravo M, Ardavin C. Monocyte-derived dendritic cells formed at the infection site control the induction of protective T helper 1 responses against Leishmania. Immunity. 2007;26(4):519–31. Epub 2007/04/07. doi: 10.1016/j.immuni.2007.01.017. PubMed PMID: 17412618.

13. Montes de Oca M, Engwerda CR, Kaye PM. Cytokines and splenic remodelling during Leishmania donovani infection. Cytokine X. 2020;2(4):100036. Epub 2021/02/20. doi: 10.1016/j.cytox.2020.100036. PubMed PMID: 33604560; PubMed Central PMCID: PMCPMC7885873.

14. Yurdakul P, Dalton J, Beattie L, Brown N, Erguven S, Maroof A, et al. Compartment-specific remodeling of splenic micro-architecture during experimental visceral leishmaniasis. Am J Pathol. 2011;179(1):23–9. Epub 2011/06/28. doi: 10.1016/j.ajpath.2011.03.009. PubMed PMID: 21703391; PubMed Central PMCID: PMCPMC3123882.

15. Abidin BM, Hammami A, Stager S, Heinonen KM. Infection-adapted emergency hematopoiesis promotes visceral leishmaniasis. PLoS Pathog. 2017;13(8):e1006422. Epub 2017/08/09. doi: 10.1371/journal.ppat.1006422. PubMed PMID: 28787450; PubMed Central PMCID: PMCPMC5560750.

16. Pinto AI, Brown N, Preham O, Doehl JSP, Ashwin H, Kaye PM. TNF signalling drives expansion of bone marrow CD4+ T cells responsible for HSC exhaustion in experimental visceral leishmaniasis. PLoS Pathog. 2017;13(7):e1006465. Epub 2017/07/04. doi: 10.1371/journal.ppat.1006465. PubMed PMID: 28671989; PubMed Central PMCID: PMCPMC5510901.

17. Preham O, Pinho FA, Pinto AI, Rani GF, Brown N, Hitchcock IS, et al. CD4(+) T Cells Alter the Stromal Microenvironment and Repress Medullary Erythropoiesis in Murine Visceral Leishmaniasis. Front Immunol. 2018;9:2958. Epub 2019/01/09. doi: 10.3389/fimmu.2018.02958. PubMed PMID: 30619317; PubMed Central PMCID: PMCPMC6305626.

18. Moore KW, O’Garra A, de Waal Malefyt R, Vieira P, Mosmann TR. Interleukin-10. Annu Rev Immunol. 1993;11:165–90. Epub 1993/01/01. doi: 10.1146/annurev.iy.11.040193.001121. PubMed PMID: 8386517.

19. Murphy ML, Wille U, Villegas EN, Hunter CA, Farrell JP. IL-10 mediates susceptibility to Leishmania donovani infection. Eur J Immunol. 2001;31(10):2848–56. Epub 2001/10/10. doi: 10.1002/1521-4141(2001010)31:10<2848::aid-immu2848>3.0.co;2-t. PubMed PMID: 11592059.

20. Stager S, Maroof A, Zubairi S, Sanos SL, Kopf M, Kaye PM. Distinct roles for IL-6 and IL-12p40 in mediating protection against Leishmania donovani and the expansion of IL-10+ CD4+ T cells. Eur J Immunol. 2006;36(7):1764–71. Epub 2006/06/23. doi: 10.1002/eji.200635937. PubMed PMID: 16791879; PubMed Central PMCID: PMCPMC2659577.

21. Owens BM, Beattie L, Moore JW, Brown N, Mann JL, Dalton JE, et al. IL-10-producing Th1 cells and disease progression are regulated by distinct CD11c(+) cell populations during visceral leishmaniasis. PLoS Pathog. 2012;8(7):e1002827. Epub 2012/08/23. doi: 10.1371/journal.ppat.1002827. PubMed PMID: 22911108; PubMed Central PMCID: PMCPMC3406093.

22. Maroof A, Beattie L, Zubairi S, Svensson M, Stager S, Kaye PM. Posttranscriptional regulation of II10 gene expression allows natural killer cells to express immunoregulatory function. Immunity. 2008;29(2):295–305. Epub 2008/08/15. doi: 10.1016/j.immuni.2008.06.012. PubMed PMID: 18701085; PubMed Central PMCID: PMCPMC2656759.

23. Chandra D, Naik S. Leishmania donovani infection down-regulates TLR2-stimulated IL-12p40 and activates IL-10 in cells of macrophage/monocytic lineage by modulating MAPK pathways through a contact-dependent mechanism. Clin Exp Immunol. 2008;154(2):224–34. Epub 2008/09/10. doi: 10.1111/j.1365-2249.2008.03741.x. PubMed PMID: 18778366; PubMed Central PMCID: PMCPMC2612710.

24. Kong F, Saldarriaga OA, Spratt H, Osorio EY, Travi BL, Luxon BA, et al. Transcriptional Profiling in Experimental Visceral Leishmaniasis Reveals a Broad Splenic Inflammatory Environment that Conditions Macrophages toward a Disease-Promoting Phenotype. PLoS Pathog. 2017;13(1):e1006165. Epub 2017/02/01. doi: 10.1371/journal.ppat.1006165. PubMed PMID: 28141856; PubMed Central PMCID: PMCPMC5283737.

25. Parmar N, Chandrakar P, Kar S. Leishmania donovani Subverts Host Immune Response by Epigenetic Reprogramming of Macrophage M(Lipopolysaccharides + IFN-gamma)/M(IL-10) Polarization. J Immunol. 2020;204(10):2762–78. Epub 2020/04/12. doi: 10.4049/jimmunol.1900251. PubMed PMID: 32277055.

26. Roy S, Mukhopadhyay D, Mukherjee S, Moulik S, Chatterji S, Brahme N, et al. An IL-10 dominant polarization of monocytes is a feature of Indian Visceral Leishmaniasis. Parasite Immunol. 2018;40(7):e12535. Epub 2018/05/11. doi: 10.1111/pim.12535. PubMed PMID: 29745990.

27. Silva-Barrios S, Smans M, Duerr CU, Qureshi ST, Fritz JH, Descoteaux A, et al. Innate Immune B Cell Activation by Leishmania donovani Exacerbates Disease and Mediates Hypergammaglobulinemia. Cell Rep. 2016;15(11):2427–37. Epub 2016/06/07. doi: 10.1016/j.celrep.2016.05.028. PubMed PMID: 27264176.

28. Meng L, Almeida LN, Clauder AK, Lindemann T, Luther J, Link C, et al. Bone Marrow Plasma Cells Modulate Local Myeloid-Lineage Differentiation via IL-10. Front Immunol. 2019;10:1183. Epub 2019/06/20. doi: 10.3389/fimmu.2019.01183. PubMed PMID: 31214168; PubMed Central PMCID: PMCPMC6555095.

29. Terrazas C, Varikuti S, Oghumu S, Steinkamp HM, Ardic N, Kimble J, et al. Ly6C(hi) inflammatory monocytes promote susceptibility to Leishmania donovani infection. Sci Rep. 2017;7(1):14693. Epub 2017/11/02. doi: 10.1038/s41598-017-14935-3. PubMed PMID: 29089636; PubMed Central PMCID: PMCPMC5665970.

30. Sato N, Kuziel WA, Melby PC, Reddick RL, Kostecki V, Zhao W, et al. Defects in the generation of IFN-gamma are overcome to control infection with Leishmania donovani in CC chemokine receptor (CCR) 5-, macrophage inflammatory protein-1 alpha-, or CCR2-deficient mice. J Immunol. 1999;163(10):5519–25. Epub 1999/11/24. PubMed PMID: 10553079.

31. MacKinnon AC, Farnworth SL, Hodkinson PS, Henderson NC, Atkinson KM, Leffler H, et al. Regulation of alternative macrophage activation by galectin-3. J Immunol. 2008;180(4):2650–8. Epub 2008/02/06. doi: 10.4049/jimmunol.180.4.2650. PubMed PMID: 18250477.

32. Reith W, Muhlethaler-Mottet A, Masternak K, Villard J, Mach B. The molecular basis of MHC class II deficiency and transcriptional control of MHC class II gene expression. Microbes Infect. 1999;1(11):839–46. Epub 1999/12/30. doi: 10.1016/s1286-4579(99)00235-x. PubMed PMID: 10614000.

33. Teh YC, Ding JL, Ng LG, Chong SZ. Capturing the Fantastic Voyage of Monocytes Through Time and Space. Front Immunol. 2019;10:834. Epub 2019/05/02. doi: 10.3389/fimmu.2019.00834. PubMed PMID: 31040854; PubMed Central PMCID: PMCPMC6476989.

34. Nguyen Hoang AT, Liu H, Juarez J, Aziz N, Kaye PM, Svensson M. Stromal cell-derived CXCL12 and CCL8 cooperate to support increased development of regulatory dendritic cells following Leishmania infection. J Immunol. 2010;185(4):2360–71. Epub 2010/07/14. doi: 10.4049/jimmunol.0903673. PubMed PMID: 20624948.

35. Svensson M, Kaye PM. Stromal-cell regulation of dendritic-cell differentiation and function. Trends Immunol. 2006;27(12):580–7. Epub 2006/10/20. doi: 10.1016/j.it.2006.10.006. PubMed PMID: 17049923.

36. Ashwin H, Seifert K, Forrester S, Brown N, MacDonald S, James S, et al. Tissue and host species-specific transcriptional changes in models of experimental visceral leishmaniasis. Wellcome Open Res. 2018;3:135. Epub 2019/01/11. doi: 10.12688/wellcomeopenres.14867.2. PubMed PMID: 30542664; PubMed Central PMCID: PMCPMC6248268.

37. Forrester S, Siefert K, Ashwin H, Brown N, Zelmar A, James S, et al. Tissue-specific transcriptomic changes associated with AmBisome(R) treatment of BALB/c mice with experimental visceral leishmaniasis. Wellcome Open Res. 2019;4:198. Epub 2020/01/25. doi: 10.12688/wellcomeopenres.15606.1. PubMed PMID: 31976381; PubMed Central PMCID: PMCPMC6961418.

38. Ohmura-Hoshino M, Goto E, Matsuki Y, Aoki M, Mito M, Uematsu M, et al. A novel family of membrane-bound E3 ubiquitin ligases. J Biochem. 2006;140(2):147–54. Epub 2006/09/07. doi: 10.1093/jb/mvj160. PubMed PMID: 16954532.

39. Thibodeau J, Bourgeois-Daigneault MC, Huppe G, Tremblay J, Aumont A, Houde M, et al. Interleukin-10-induced MARCH1 mediates intracellular sequestration of MHC class II in monocytes. Eur J Immunol. 2008;38(5):1225–30. Epub 2008/04/05. doi: 10.1002/eji.200737902. PubMed PMID: 18389477; PubMed Central PMCID: PMCPMC2759377.

40. Mittal SK, Cho KJ, Ishido S, Roche PA. Interleukin 10 (IL-10)-mediated Immunosuppression: MARCH-I INDUCTION REGULATES ANTIGEN PRESENTATION BY MACROPHAGES BUT NOT DENDRITIC CELLS. J Biol Chem. 2015;290(45):27158–67. Epub 2015/09/27. doi: 10.1074/jbc.M115.682708. PubMed PMID: 26408197; PubMed Central PMCID: PMCPMC4646393.

41. Galbas T, Raymond M, Sabourin A, Bourgeois-Daigneault MC, Guimont-Desrochers F, Yun TJ, et al. MARCH1 E3 Ubiquitin Ligase Dampens the Innate Inflammatory Response by Modulating Monocyte Functions in Mice. J Immunol. 2017;198(2):852–61. Epub 2016/12/13. doi: 10.4049/jimmunol.1601168. PubMed PMID: 27940660.

