## Supplemental Figure 1 for "Interferon-γ-producing CD4^+^ T cells drive monocyte activation in the bone marrow during experimental *Leishmania donovani* infection"

### Supplementary Figure 1

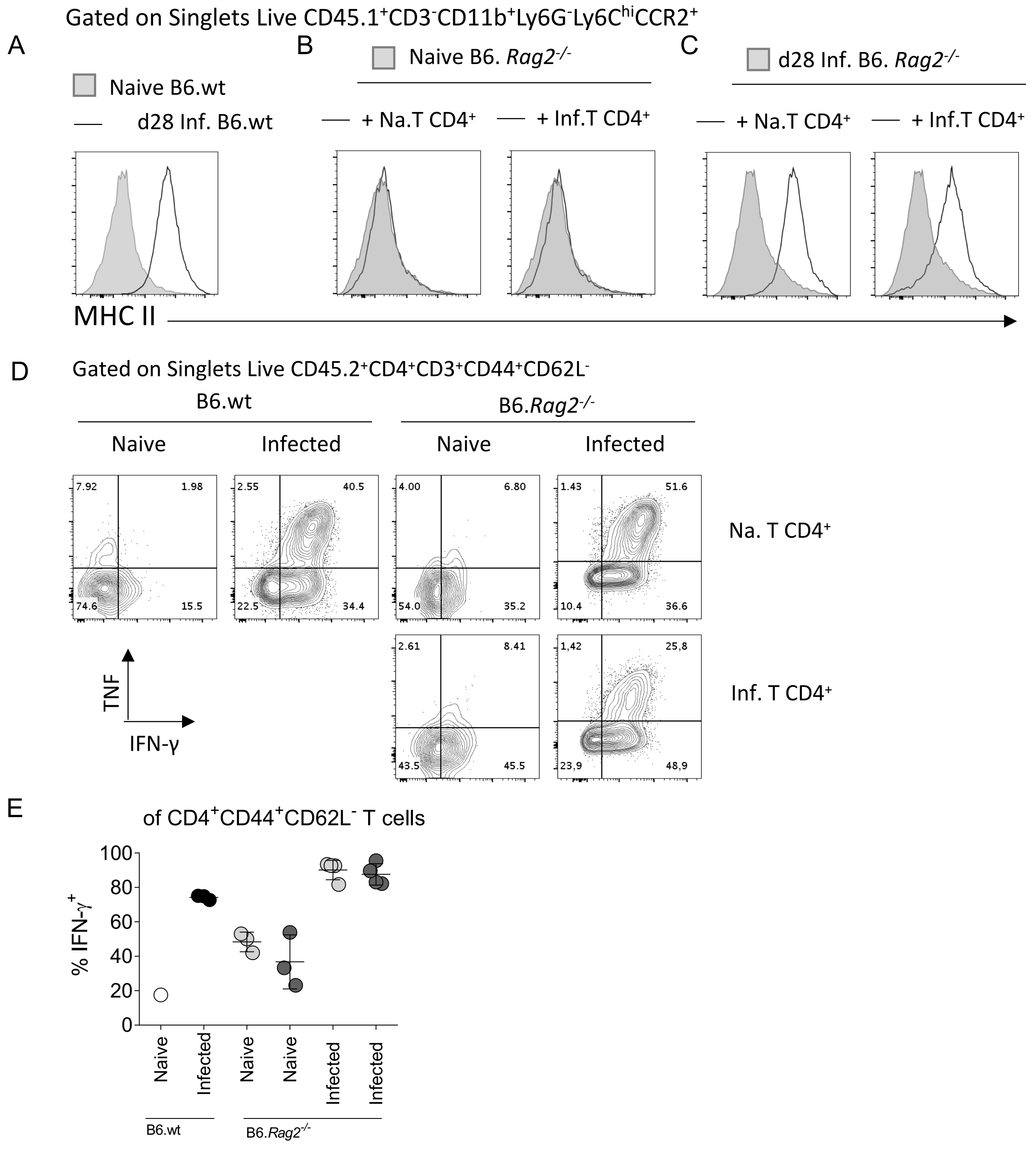


**Figure S1 : CD4 T cells fail to activate iMO in absence of *L. donovani* infection.** BM $CD4^+$ T cells from naive (Na.T $CD4^+$) or infected (Inf.T $CD4^+$) B6.wt mice were adoptively transfered to naive B6.*Rag2*$^{-/-}$ **A-C.** MHCII expression on iMo in the BM (**A**) of naive and d28 infected B6.wt mice, of naive (**B**) and d28 infected (**C**) B6.*Rag2*$^{-/-}$ 2 weeks post adoptive transfer shown as representative histogram plots. **D.** Cytokine production by adoptively transfered BM $CD4^+$ T cells 2 weeks post transfer into recipient mice shown as representative dot plots. Cells were stimulated with PMA/ionomycin. **E**. Percentage of IFNγ produced by BM $CD4^+$ T cells. Data are derived from analysis of 3 to 5 individual mice of each strain per group and are shown as mean ± SD. Data are representative of 2 independant experiments.
