## Supplemental Figure 2 for "Interferon-γ-producing CD4^+^ T cells drive monocyte activation in the bone marrow during experimental *Leishmania donovani* infection"

**Supplementary Figure 2**

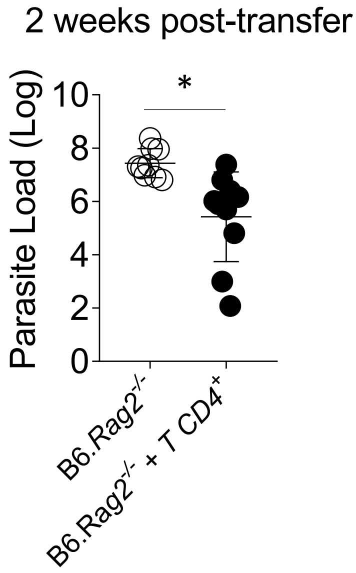


**Figure S2 : Adoptive transfer of BM CD4 T cells reduces parasite load in B6.*Rag2*$^{-/-}$ mice** Parasite load per one million cells in the bone marrow measured by limiting dilution assay. Data are derived from analysis of 8 and 10 individual mice in B6.*Rag2*$^{-/-}$ group and in B6.*Rag2*$^{-/-}$+ T CD4$^{+}$ cells respectively. Data are pooled from 3 independent experiments at 2 weeks post transfer are shown as mean ± SD. *, $p<0.05$
