## Supplemental Figure 3 for "Interferon-γ-producing CD4^+^ T cells drive monocyte activation in the bone marrow during experimental *Leishmania donovani* infection"

**Supplementary Figure 3**

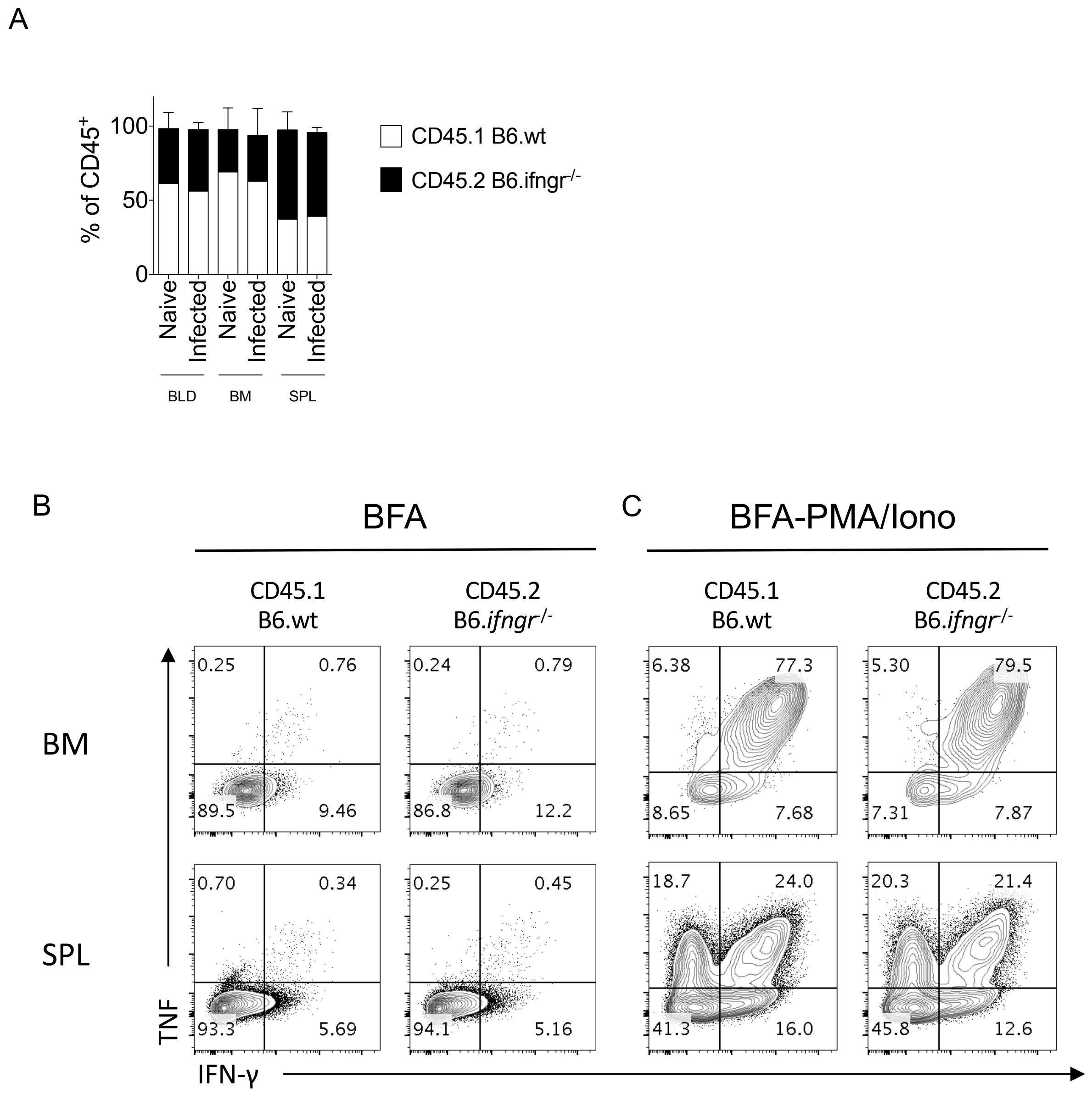


**Figure S3 : Cytokine production from CD4+ T cells in mixed bone marrow chimeras**

Naïve lethally irradiated B6.CD45.1 recipient mice received a 50:50 of BM cells from B6.CD45.1 and B6.*ifngr1*$^{-/-}$ CD45.2 mice. Mice were subsequently infected with $3x10^7$ *L. donovani* amastigotes for 28 days. **A**. Percentage of CD45.1 and CD45.2 of singlets live CD45$^+$ cell population in the blood, BM and spleen. **B-C**. Cytokine production by CD4$^+$ T cells in the BM and the spleen. Cells were stimulated with (**B**) BFA alone or (**C**) PMA/Ionomycin. Data are the pool of 5 mice/group/experiment. Mean +/- SD is shown.
